# Sequence Characterization of Glutamate Receptor Genes of Rat a Vertebrate and Arabidopsis Thaliana a Plant

**DOI:** 10.1101/273334

**Authors:** Antara Sengupta, Pabitra Pal Choudhury, Subhadip Chakraborty

**Affiliations:** Dept. of MCA, MCKV Institute of Engineering, Howrah, India; Applied Statistics Unit, Indian Statistical Institute, Kolkata, India; Dept. of Botany, Nabadwip Vidyasagar College, Nabadwip, India

**Keywords:** Glutamate Receptors, Arabidopsis thaliana, Chemical Properties, Directed Graph, Phylogenetic Tree

## Abstract

iGluR gene family of a vertebrate, Rat and AtGLR gene family of a plant, Arabidopsis thaliana [4] perform some common functionalities in neuro-transmission, which have been compared quantitatively. Our attempt is based on the chemical properties of amino acids [6, 7, 8] comprising the primary protein sequences of the aforesaid genes. 19 AtGLR genes of length varying from 808 amino acid (aa) to 1039 aa and 16 iGluR genes length varying from 902aa to 1482 aa have been taken as data sets. Thus, we detected the commonalities (conserved elements) during the long evolution of plants and animals from a common ancestor [4]. Eight different conserved regions have been found based on individual amino acids. Two different conserved regions are also found, which are based on chemical groups of amino acids. We have tried too to find different possible patterns which are common throughout the data set taken. 9 such patterns have been found with size varying from 2 to 5 amino acids at different regions in each primary protein sequences. Phylogenetic trees of AtGLR and iGluR families have also been constructed. This approach is likely to shed light on the long course of evolution.

## I. INTRODUCTION

Plants grow in silence but react to wounding. Specific genes are there in plants which make control over such functionalities in neuro-transmission. On the other hand, animals have well developed nervous systems. Mammals for example can relay electrical signals at a high speed of 100 m/sec approximately [1]. Earlier it was believed that plants do not have anything like a nervous system, they do not respond to stimuli in the way animals do. But according to the great pioneer Scientist Sir Acharya J.C.Bose,[3] plant also has an alternative sort of sensitive nervous system, not like that of animals but can responds to various external stimuli. Numerous works since Darwin [5] have proved the response of plants to various signals like light, touch, wounding, stimulating but now it has been reported that glutamate receptors of vertebrate nervous system and similar receptors in the plant equivalent of nerve have so close similarities that they must have evolved from a common ancestor [4]. Arabidopsis genome sequencing has discovered a large family of Glutamate Receptor like genes (AtGLR) in them. It has been reported that like ionotropic glutamate receptor (iGluR) gene family of animals, AtGLR gene family in Arabidopsis have some functions in neurotransmission. (GLRs) in Arabidopsis thaliana are organism those contribute in neurotransmission in it. Glutamate receptors are also expressed by nonneuronal cells, including neuroglia and T lymphocytes, where, as in neurons, they serve to convey glutamate signals across the plasma membrane. Ionotropic glutamate receptors are essential for mammalian Central Nervous System. They intervene in chemical synaptic transmission at the clear majority of excitatory synapses, underlie well-characterized cellular models of learning and memory, modulate excitability of neuronal networks. KA1 mRNA (also known as GRIK4) has functionality in only CA3 of hippocampus, whereas, KA2 mRNA (also known as GRIK4) can be detected almost all region of rat brain [22]. Mutation in both genes can lead to neurological and psychiatric disorder. Ionotropic Glutamate receptor, delta-1 and delta-2 are the subunits, which extend the excitatory amino acid receptor family of species rat. Delta-1 is expressed in the caudate putamen and delta-2 is expressed predominantly in Purkinje cells of the cerebellum [23].Amino acids are organic compounds containing specific biochemical compositions that instruct primary protein sequences to have specific structure, to perform specific functions and to get stability. Quantitative understanding of genes at molecular level requires some Mathematical parameters to evaluate some data, to derive some values and to verify some facts. Similarity/Dissimilarity analysis is one of the fundamental methods used to investigate evolutionary relationship between various species [6].

Analysis of DNA as well as amino acid sequences using underlying biological information hidden in them is an old approach, whereas Graphical representation of primary protein sequence give alignment free method which provides visual inspection of the sequence and instant provision of analyzing similarities with others. Numerous authors [8-25] proposed several 2D and 3D graphical representations of DNA as well as amino acid sequences.. Some research papers are reported, where multiplet structure of amino acid sequences are the theme to investigate evolutionary relationship among the species [22]. Attempt has also been made to find out invariant conservation of certain amino acids of various organism using biochemical properties of amino acid side chains [6-8].Several works have also been done with PPIs, where it is tried to find whether hub and the genes in network exhibit any kind of proximity with respect to the codons[27].

In this paper, the section of materials and methods has several sub sections. In first subsection firstly, the classifications of the 20 amino acids have been taken place on the basis of their chemical nature. As our primary objective is to make quantitative comparison between iGluR gene family of mammal (Rat) and AtGLR of plant (Arabidopsis thaliana) based on their chemical features and investigate evolutionary relationships among them, so in this context, the data sets on which the experiments are carried out has been specified in the second sub section. In third sub section the methodologies have been described thoroughly. In third section, the procedures stated are also applied on two data sets taken and experimental results are discussed clearly. Finally, total work has been concluded in the last section. Thus, the novelty lies in the mathematical modeling to establish the fact that like Glutamate Receptor gene family in mammal, plant also has Glutamate Receptors like genes which have been conserved during the long process of their evolution from a common ancestry.

## II. METHODOLOGY

### A. Classification of Amino Acids

The chemical features of 20 amino acids are basically depended on eight chemical properties of them. A protein structure is dependent upon the chemical properties of amino acid by which it is formed. Hence it determines the biological activity of proteins. It has previously been established that amino acid properties are not only decisive in determining protein structure and functions performed by them, but also play a role in molecular evolution [6,7].

To understand the structural and chemical property of Amino Acid one need to understand the structural and chemical property of protein. It has been reported in a paper that molecular evolutionary analysis of conserved region based on chemical properties of the side chain of amino acids can determine the essential amino acids in the core catalytic region [8]. So here in this manuscript it is aimed to understand how the biochemical nature of each amino acid sequences figure out evolutionary conserved regions among glutamate genes of mammal and plant during the long process of evolution. Analyze molecular evolutionary carry out the further experiments based on 8 distinct chemical properties of amino acids. So, 20 amino acids are clustered or classified into 8 groups accordingly as shown in Table 1. Eight different numeric values are assigned to each group.

**Table 1:** Classification of amino acids as per their Chemical Properties and numeric representation of each Class.

| Group Name | Amino Acids included | Group/Class No. |
| --- | --- | --- |
| Acidic | Aspartate (D), Glutamate (E) | 1 |
| Basic | Arginine (R), Histidine (H), Lysine (K) | 2 |
| Aromatic side chain | Tyrosine (Y), Phenylalanine (F), Tryptophan (W) | 3 |
| Aliphatic side chain | Isoleucine (I), Leucine (L), Valine (V), Alanine (A), Glycine (G) | 4 |
| Cyclic | Proline (P) | 5 |
| Sulfur containing | Methionine (M), Cysteine (C) | 6 |
| Hydroxyl containing | Serine (S), Threonine (T) | 7 |
| Acidic amide | Glutamine (Q), Asparagine (N) | 8 |

### B. Data Set Specification

To carry out the entire experiment, AtGLR gene family of Arabidopsis thaliana and Glutamate receptor genes such as iGluR genes of Rat have been taken in account. A large family of GLR genes has been uncovered and reported in NCBI. 19 of such sequences have been selected to carry out the investigations (Table 2). 16 amino acid sequences of iGluR family of Rat have also been taken for studies which are shown in Tables 3.

**Table 2.** Data set specification of AtGLR gene family in Arabidopsis.

| Gene Name | Accession No. | Length of AA | Gene Name | Accession No. | Length of AA |
| --- | --- | --- | --- | --- | --- |
| AtGLR1.1 | AAF26802.1 | 938 aa | AtGLR2.6 | CAB96653.1 | 925 aa |
| AtGLR1.2 | BAA96960.1 | 920 aa | AtGLR2.7 | AAC33239.1 | 925 aa |
| AtGLR1.3 | BAA96961.2 | 895 aa | AtGLR2.8 | AAC33237.1 | 962 aa |
| AtGLR1.4 | AAF02156.1 | 898 aa | AtGLR2.9 | AAC33236.1 | 895 aa |
| AtGLR2.1 | AAB61068.1 | 829 aa | AtGLR3.2 | CAA18740.1 | 1039 aa |
| AtGLR2.2 | AAD26895.1 | 906 aa | AtGLR3.3 | AAG51316.1 | 921 aa |
| AtGLR2.3 | AAD26894.1 | 934 aa | AtGLR3.4 | AAB71458.1 | 808 aa |
| AtGLR2.4 | CAA19752.1 | 958 aa | AtGLR3.5 | AAC69939.1 | 867 aa |
| AtGLR2.5 | CAB96656.1 | 940 aa | AtGLR3.6 | CAB63012.1 | 860 aa |
|  |  |  | AtGLR3.7 | AAC69938.1 | 858 aa |

**Table 3.** Data set specification of iGluR gene family in Rat.

| Gene Name | Accession No. | Length of AA | Gene Name | Accession No. | Length of AA |
| --- | --- | --- | --- | --- | --- |
| GluRA | P19490 | 1009 aa | KA-2 | Q63273 | 902 aa |
| GluRB | P19491 | 1007 aa | Delta 1 | Q62640 | 956 aa |
| GluRC | AAA41245.1 | 920 aa | Delta 2 | Q63226 | 979 aa |
| GluRD | AAA41246.1 | 908 aa | NMDA Zeta1 | P35439 | 1237 aa |
| GluR5 | AAA02873.1 | 888 aa | NMDA2A | AAC03565.1 | 938 aa |
| GluR6 | P42260 | 907 aa | NMDA2B | AAA41714.1 | 1464 aa |
| GluR7 | AAC80577.1 | 883 aa | NMDA2C | AAA41713.1 | 1482 aa |
| KA-1 | AAA17830.1 | 888 aa | NMDA(N R2D) | AAC37647.1 | 1323 aa |

### C. Numerical Representation of Data Set taken as per Classification

To characterize a primary protein sequence, it is necessary to replace each amino acid of that primary protein sequence by the corresponding numerical value of the class or group from which it belongs to. Let *S* = *S*_1_, *S*_2_,…. *S*_*k*_, be an arbitrary primary protein sequence, where for any S represents a single amino acid from the set S, such that *S*_*i*_∈ {A,C,D,E,F,G,H,I,K,L,M,N,P,Q,R,S,T,V,W,Y}.

Let 20 amino acids are classified into a set of classes C. Then, it is possible to read each amino acid of set S by its corresponding class C, where C ∈ {1,2,3,4,5,6,7,8}.As an example, according to Table1, a chunk of amino acid sequence ‘MEALT’ will be represented as ‘61447’ after the encoding into the numeric representation.

### D. Calculate Percent wise presence of each Amino Acid present in each sequence and each Group of Amino Acid in each sequence

Presence of all 20 amino acid sequences are not same for all species. Even this rate of frequency vary sequence to sequence. In this paper rate of occurrence i.e., frequency of every individual amino acid. The chemical properties of the amino acids determine the biological activity of the protein. Proteins not only act as catalyst in most of the reactions in living cells, but they control virtually all cellular processes too. In addition, amino acid sequence of a protein contains the necessary information to determine how that protein will fold into a three-dimensional structure, and the stability of the resulting structure. Sometimes the amino acid encodes some information which are inherited from the previous stage (before translation). Basic and acidic Amino Acids are essential in the formation of Beta sheets, which are secondary protein structures that are stabilized by the creation of hydrogen bonds between acidic and basic amino acids [23]. Whereas, aromatic amino acids provide a negative electrostatic potential surface that leads to cation-pi interaction. Cation -p interactions have significant contribution in overall stability of proteins. So, it is important to calculate the presence of each group of amino acid which contains specific chemical properties. In this part of the paper percentage of each group of amino acid present in each sequence has been calculated to get a quantitative understanding about the presence of each chemical group in each amino acid sequence.

### E. Common patterns/blocks finding and identifying conserved regions

Conservation of DNA or protein sequences across species indicates that a sequence has been maintained by evolution; hence conserved protein sequence regions are extremely useful to investigate, identify and study functional and structural importance of those regions. Here in this part of the manuscript firstly conserved regions are detected. Given an input of pattern length L (2<L<8), all possible combination of patterns of length L using the numerical values 1-8 are generated. Every possible pattern is investigated among the sequences, and if the pattern is found for all the sequences, the pattern is selected and stored along with the locations, otherwise the pattern is discarded. Thus, by varying pattern length L, it is possible to find patterns of different lengths along with their corresponding location.

### F. Investigating evolutionary relationships between two species

Let S= {S_1_, S_2_,…S_n_} be a set of Amino acid sequences, where each sequence S_i_ for i={1,2,…n} can be represented through weighted directed multi graph G_m_. G_m_={A,V} is say a multigraph, where set of vertices V={V_1_, V_2_,V_3_,V_4_,V_5_,V_6_,V_7_,V_8_} represent the elements of class C, where C={1,2,3,4,5,6,7,8} and arcs A represent the possible parallel arcs present between any two vertices. For any Amino acid sequence, say V_i_ and V_j_ are any two vertices for which directed multi graphs may have parallel arcs from V_i_ to V_j_. now define the weight of each arc from V_i_ to V_j_ as 1, so that for every arc from V_i_ to V_j_, total weight of the arcs of graph G_m_ is,

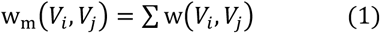

In this way the graphical representation of Amino Acid (aa) sequence can give us an idea about the distribution of chemical properties in them.Now the directed multi graph can be drawn through adjacency matrix. We can get its corresponding (8X8) adjacent matrix M for each pair of vertices as mentioned below in Table 10, which gives us 64 dimensional vectors in row order say R, where R=(w(1,1),w(1,2),…..w(1,8),, w(8,1),…w(8,8)).

**Table 4.** Representing weighted directed multigraph through adjacency matrix.

| Weight | 1 | 2 | 3 | 4 | 5 | 6 | 7 | 8 |
| --- | --- | --- | --- | --- | --- | --- | --- | --- |
| 1 | w(1,1) | w(1,2) | w(1,3) | w(1,4) | w(1,5) | w(1,6) | w(1,7) | w(1,8) |
| 2 | w(2,1) | w(2,2) | w(2,3) | w(2,4) | w(2,5) | w(2,6) | w(2,7) | w(2,8) |
| 3 | w(3,1) | w(3,2) | w(3,3) | w(3,4) | w(3,5) | w(3,6) | w(3,7) | w(3,8) |
| 4 | w(4,1) | w(4,2) | w(4,3) | w(4,4) | w(4,5) | w(4,6) | w(4,7) | w(4,8) |
| 5 | w(5,1) | w(5,2) | w(5,3) | w(5,4) | w(5,5) | w(5,6) | w(5,7) | w(5,8) |
| 6 | w(6,1) | w(6,2) | w(6,3) | w(6,4) | w(6,5) | w(6,6) | w(6,7) | w(6,8) |
| 7 | w(7,1) | w(7,2) | w(7,3) | w(7,4) | w(7,5) | w(7,6) | w(7,7) | w(7,8) |
| 8 | w(8,1) | w(8,2) | w(8,3) | w(8,4) | w(8,5) | w(8,6) | w(8,7) | w(8,8) |

Now let for any two amino acids sequences say S_1_ and S_2_ the corresponding set of vectors are P={P_1_, P_2_,…,P_64_} and Q={Q_1_, Q_2_, ….Q_64_} respectively, then weight deviation (WD) between the two sequences S_1_ and S_2_ can be represented as stated in equation 2.

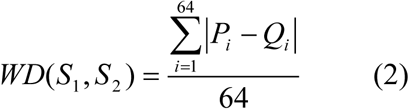

## III. APPLICATION AND RESULTS

### A. Calculate Percentage of each Group of Amino Acid present in each sequence

It has been discussed before that the chemical properties of primary protein sequences have significant role in protein folding and protein structure making. It can be observed that in both the Tables 5 and 8 group 6, i.e. amino acids of aliphatic group has remarkably major contributions to make primary protein sequences in both species, whose hydropathy profile is hydrophobic.

**Table5.**
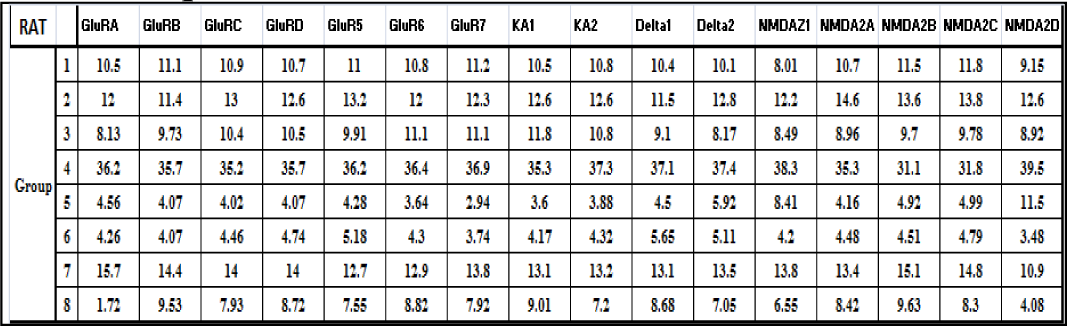
Percentage of each Group of Amino Acid present in each sequence of Rat.

**Table6.**
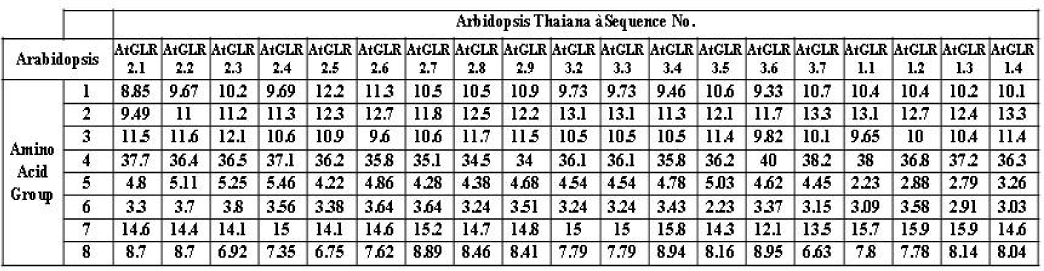
Percentage of each Group of Amino Acid present in each sequence of Arabidopsis.

**Table 7.**
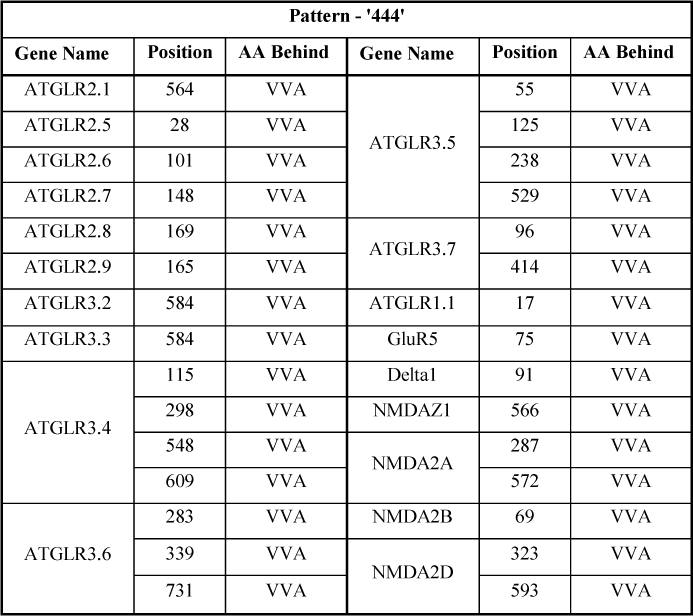
Common pattern finding (length of 3 aa) for which same amino acids are contributed in same order as per classification.

**Table 8.**
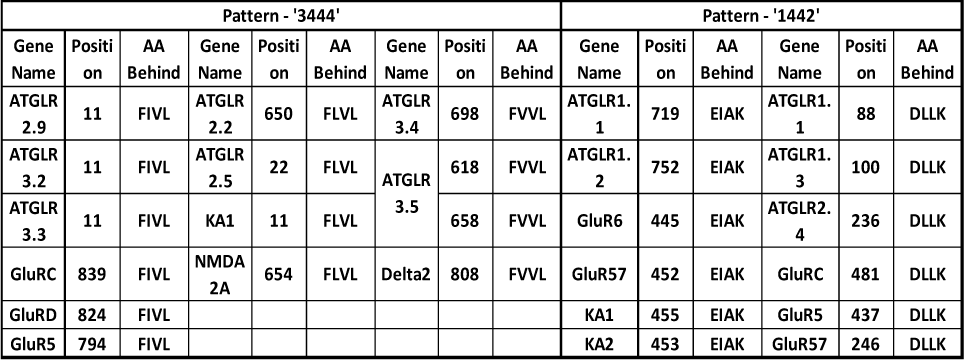
Common pattern finding (length of 4 aa) for which same amino acids are contributed in same order as per classification.

### B. Common pattern finding in all primary protein sequences

As 20 amino acids are grouped according to their chemical properties, and each of them is identified with specific numeric value, so when common pattern finding procedure has been made for all the primary protein sequences, some numerical patterns are found which are common in all the primary protein sequences taken. In this section it is tried to make microscopic view of those patterns. Table 7 contains patterns of length of 3 amino acids (aa) for which same amino acids are contributed in same order, whereas, Table 8 and 9 contain patterns of length of 4 aa and 5 aa respectively. It is remarkable that in all cases amino acids from group 4 (aliphatic) has major contribution in pattern forming. As it is known that Aliphatic R groups are hydrophobic and nonpolar, those are one of the major driving forces for protein folding, give stability to globular or binding structures of protein [23].It has been observed in Table 9 that ‘VVA’ is a block of length 3 aa which is found in majority of primary protein sequences of Arabidopsis and 6 of Rat.Table 10 shows blocks having lengths of 4 aa. ‘FIVL’, ‘FLVL’, ‘FVVL’, ‘EIAK’ and ‘DLLK’ are the 5 blocks which are found in some primary protein sequences of both the species.

**Table 9.**
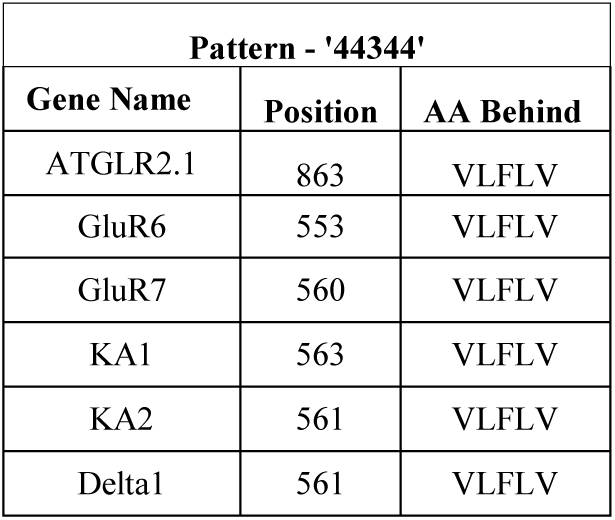
Common pattern finding (length of 5 aa) for which same amino acids are contributed in same order as per classification.

| Pattern - '44344' |  |  |
| --- | --- | --- |
| Gene Name | Position | AA Behind |
| ATGLR2.1 | 863 | VLFLV |
| GluR6 | 553 | VLFLV |
| GluR7 | 560 | VLFLV |
| KA1 | 563 | VLFLV |
| KA2 | 561 | VLFLV |
| Delta1 | 561 | VLFLV |

**Table 10.**
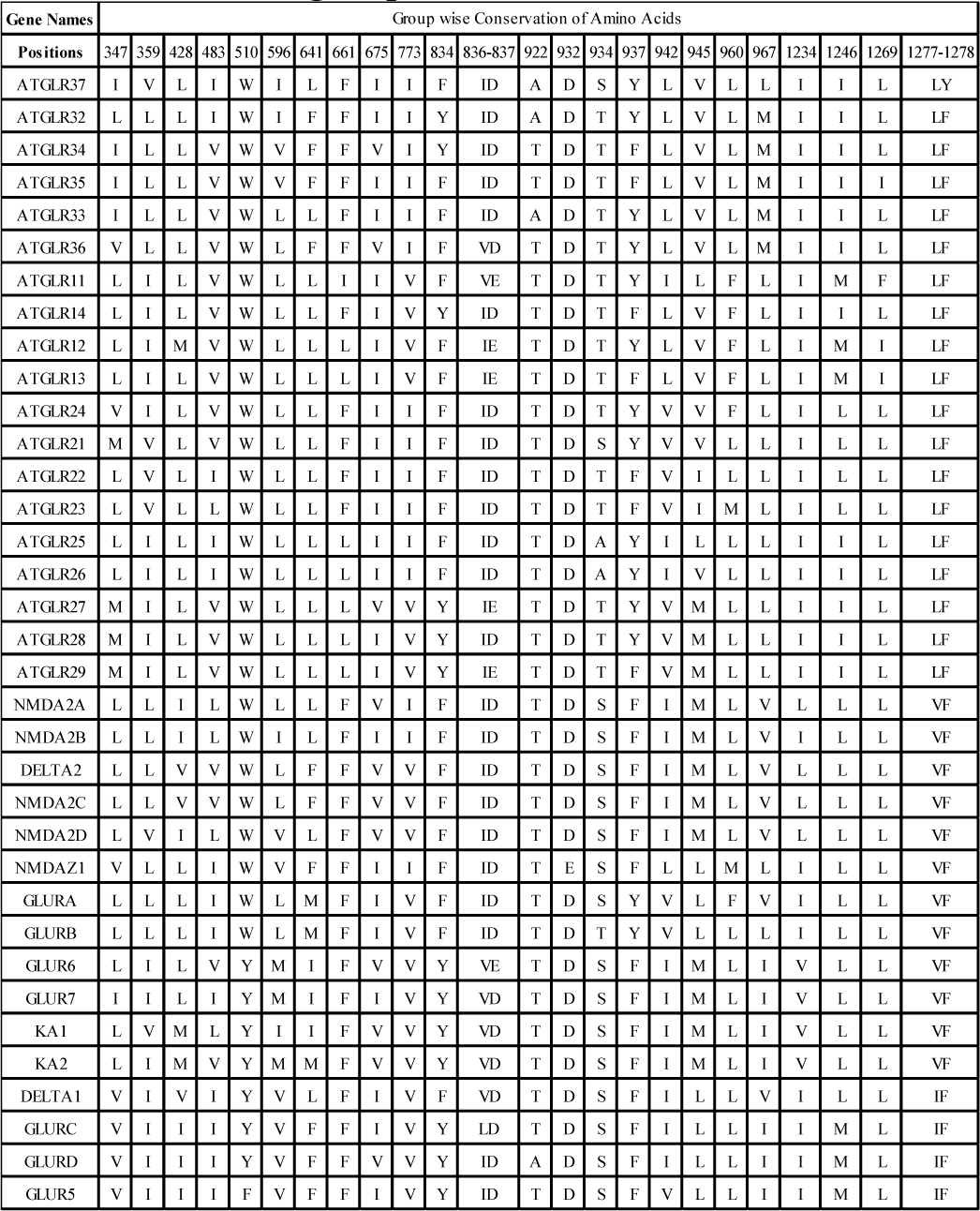
Various conserved regions of all primary protein sequences taken. The regions specified are conserved based on chemical groups of amino acids.

**Table 11.**
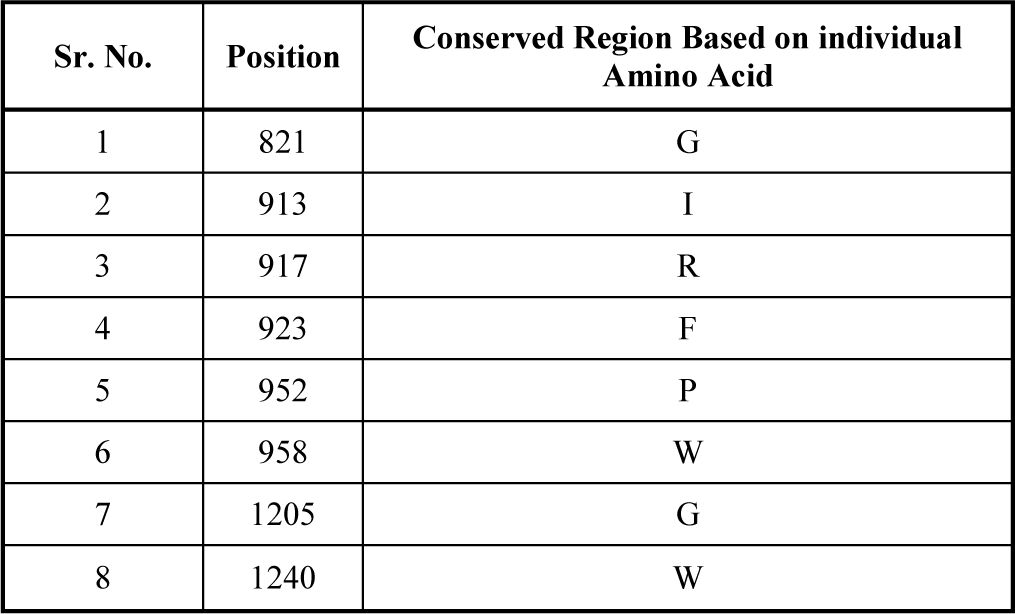
Various conserved regions of all primary protein sequences taken based on individual amino acid.

Table 9 shows common block ‘VLFLV’ of 5 aa length, which is found in AtGLR2.1 of Arabidopsis and in GluR6, GluR7, KA1, KA2, Delta1 of Rat. Table 10 shows regions in each primary protein sequences, where the regions specified are conserved based on chemical groups of amino acids. The regions are varying from 1aa to 2 aa long. It can be easily observed that the conserved regions are majorly belong to aliphatic group of chemical propertiesThe regions which are conserved based on individual amino acid are shown in the table 13 along with the regions of conservations. G, I, R, F, P, W, G and W are those amino acids. These types of conservations depict regional importance of those patterns from functional and structural point of view to make 3D protein structures of each primary protein sequence.

### C. Investigating evolutionary relationships between two species

To investigate and identify evolutionary relationships between the data sets taken, firstly equation 1 has been used to derive 8X8 weight matrix for each AA sequences. As examples weight matrices for AtGLR1.1 of Arabidopsis has been stated below in Tables 12. Equation 2 mentioned in previous section is used to get weight deviation (WD) between each pair of AA sequences taken. Thus, a distance matrix say dissimilarity matrix has been formed to analyze evolutionary relationships existing between them.

**Table12.** Weighted directed multigraph through adjacency matrix of AtGLR1.1.

| Vertex Vs. Vertex | 1 | 2 | 3 | 4 | 5 | 6 | 7 | 8 |
| --- | --- | --- | --- | --- | --- | --- | --- | --- |
| 1 | 11 | 17 | 9 | 31 | 2 | 3 | 9 | 8 |
| 2 | 20 | 10 | 16 | 41 | 1 | 1 | 13 | 7 |
| 3 | 7 | 10 | 8 | 32 | 4 | 4 | 13 | 9 |
| 4 | 30 | 37 | 33 | 122 | 8 | 8 | 59 | 22 |
| 5 | 0 | 2 | 4 | 11 | 0 | 3 | 2 | 3 |
| 6 | 3 | 4 | 4 | 12 | 1 | 2 | 4 | 1 |
| 7 | 15 | 19 | 8 | 48 | 5 | 6 | 25 | 12 |
| 8 | 4 | 11 | 5 | 22 | 4 | 3 | 13 | 5 |

Fig 2 is a phylogenetic tree, which is being constructed from the dissimilarity matrix specified in Supplementary table (ST1) attached. So, all the 16 primary protein sequences of Rat and 19 primary sequences of Arabidopsis are participated in this figure. According to fig 2, AtGLR2.1 to AtGLR 2.9 and AtGLR3.2 to AtGLR 3.7 genes of Arabidopsis Thaliana except AtGLR2.5 and AtGLR3.6 have close evolutionary relationships with GluRA and GluRB of Rat, whereas, AtGLR3.6 are close to NMDAZ1, NMDA2B, NMDA2C and NMDA2D of Rat.In fig 3 phylogenetic tree has been constructed with all the 19 AtGLR genes of Arabidopsis thaliana and 8 iGluR genes (GluRA, GluRB, GluRC, GluRD, GluR5, GluR6, GluR7, GluR8) of Rat to get more finer view about evolutionary relationships between glutamate gene families of Rat and Arabidopsis. According to the phylogenetic AtGLR1.1, AtGLR1.2, AtGLR1.3, AtGLR1.4 and AtGLR2.5 genes have similarities in chemical properties with 8 iGluR genes GluRA, GluRB, GluRC, GluRD, GluR5, GluR6, GluR7 and GluR8 of vertebrate Rat. The observation thus depicts the evolutionary relationships of ionotropic glutamate receptor (iGluR) genes of vertebrates with AtGLR genes in Arabidopsis.

**Figure1:**
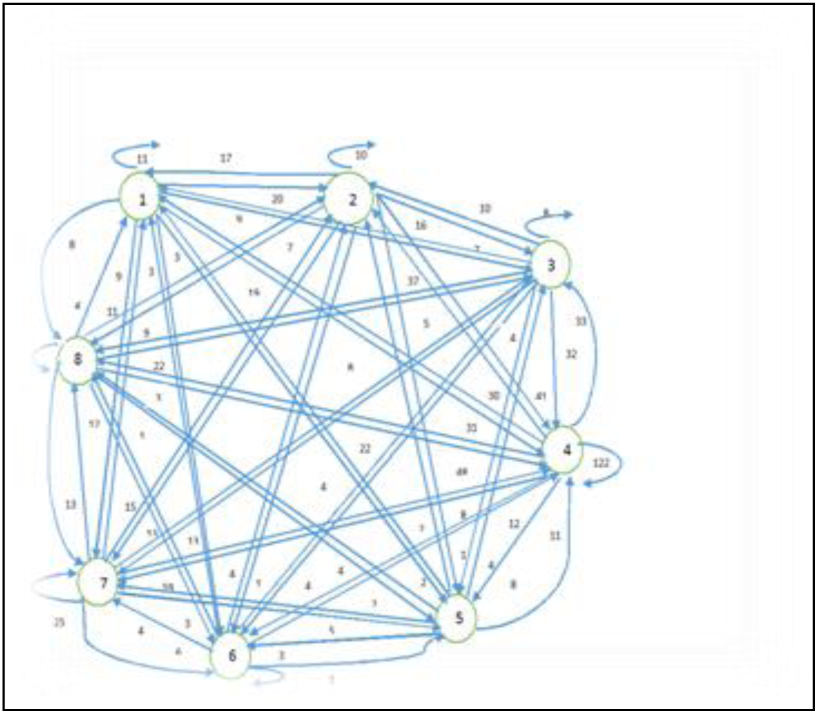
Corresponding weighted directed multi graphical representation of AtGLR1.1 has been shown as an example.

**Figure2:**
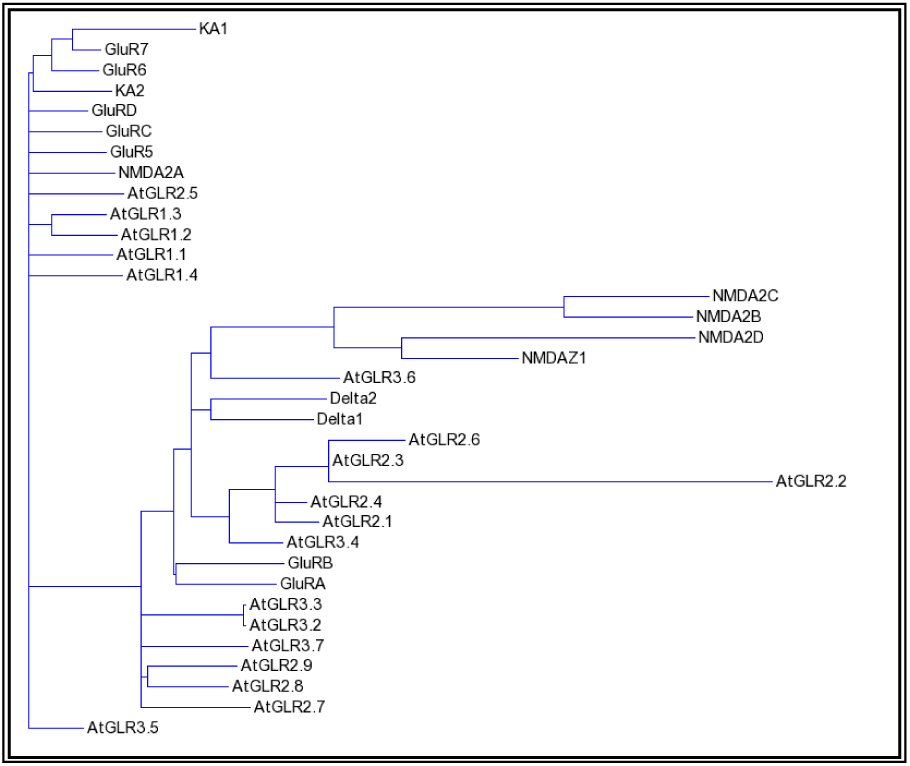
Phylogenetic tree constructed for total Data Sets taken.

**Figure 3:**
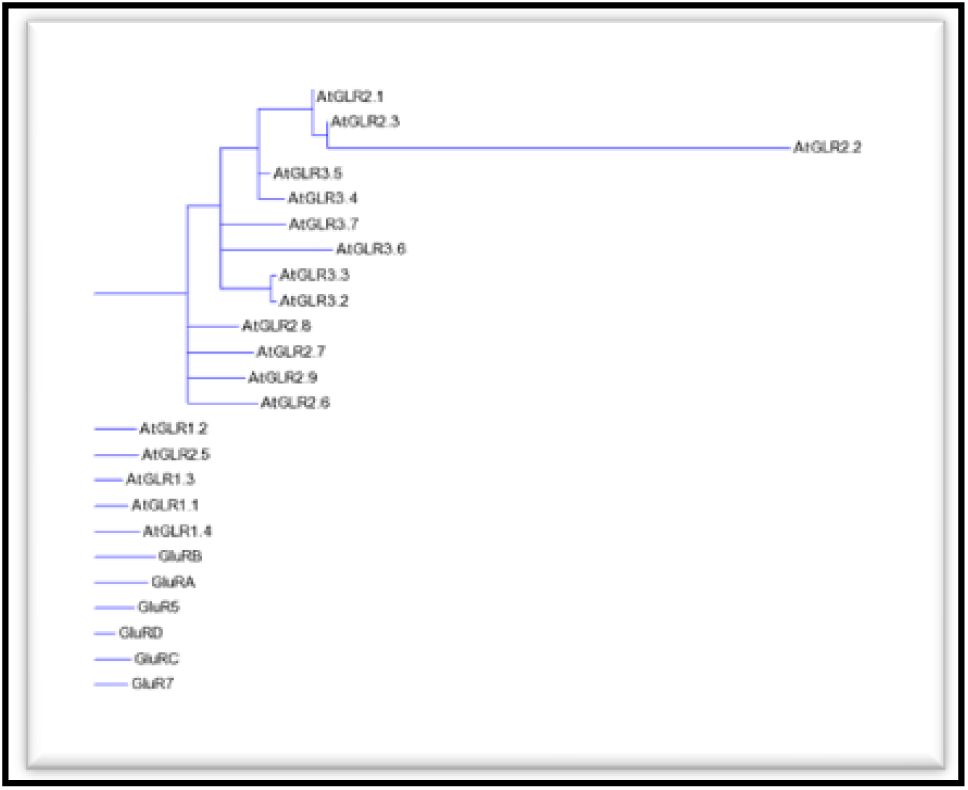
Phylogenetic tree constructed of all the 19 AtGLR genes of Arabidopsis thaliana and 8 iGluR genes (GluRA, GluRB, GluRC, GluRD, GluR5, GluR6, GluR7, GluR8)

## IV. CONCLUSION AND DISCUSSION

The analysis throughout the paper confirms in silico the recent biological claim that the iGluR family of the rat, a mammal and AtGLR family of Arabidopsis, a plant has some common functionalities in neurotransmission and have been conserved during the long process of evolution from a common ancestor. The graph theoretic approach which is applied to investigate evolutionary relationship among all the primary protein sequences taken is a completely alignment free method and hence the time complexity is directly proportional to the sequence length N, that is O(N).

## ACKNOWLEDGMENTS

The authors are grateful to Professor R.L. Brahmachary for suggesting the problem and encouraging them and also to Mr. Jayanta Kr. Das, Mr. Arindam Sengupta and Dr. Santanu Sen for their valuable suggestions.

---

**ST1:**
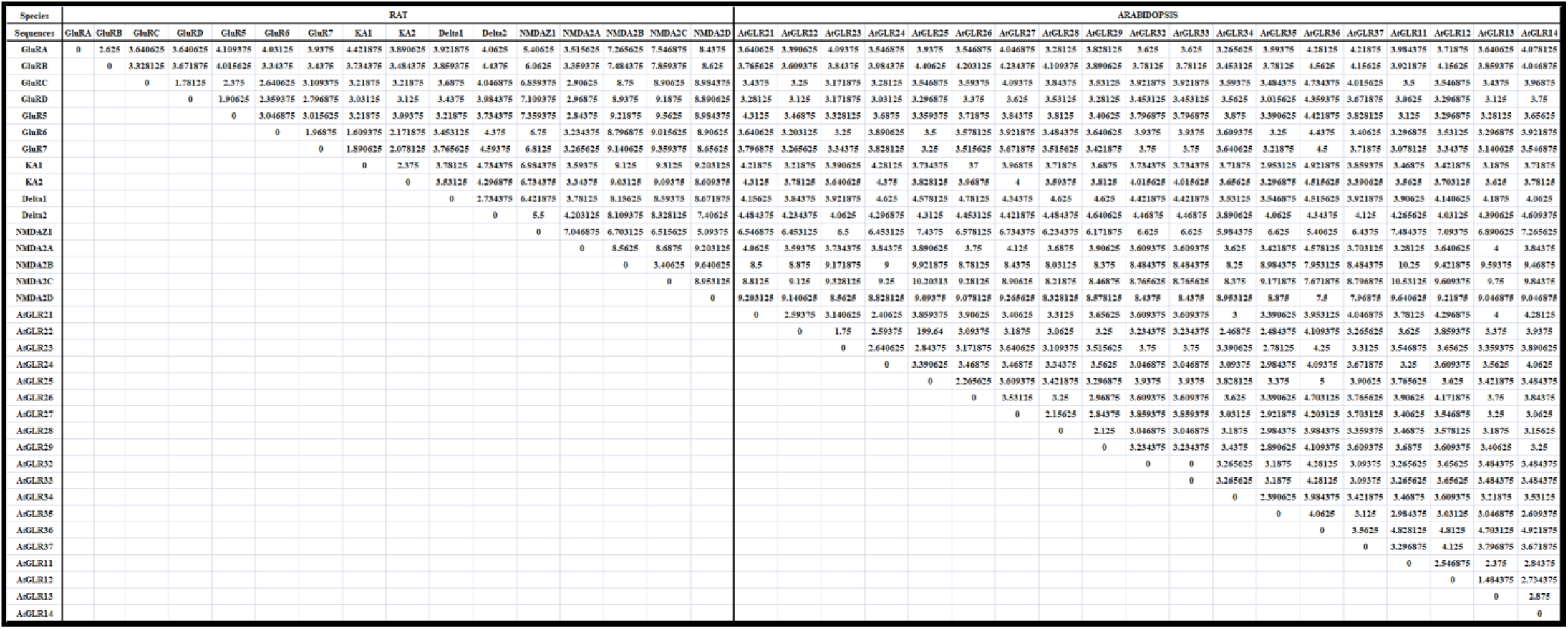
Dissimilarity matrix of all primary protein sequences taken to investigate evolutionary distance between them.

